# Characterization of lymphoma models for the surface ROR1 expression

**DOI:** 10.1101/2025.11.19.684505

**Authors:** Chiara Tarantelli, Elisa Civanelli, Filippo Spriano, Giorgia Risi, Luciano Cascione, Gunnar Kaufmann, Francesco Bertoni

## Abstract

Receptor tyrosine kinase-like orphan receptor 1 (ROR1) is a developmental antigen aberrantly expressed in several B-cell malignancies and represents an attractive therapeutic target. Here, we systematically characterized ROR1 expression in a large panel of B-cell lymphoma cell lines and assessed the activity of a ROR1-targeting antibody-drug conjugate (ADC). ROR1 RNA expression was first evaluated using a previously generated total RNA-Seq dataset from 47 B-cell lymphoma cell lines. ROR1 transcripts were detectable in most models, with 19 cell lines showing moderate to high expression. To determine whether ROR1 is present on the cell surface, we next analyzed 29 cell lines by flow cytometry using a PE-conjugated anti-ROR1 antibody. Only a subset of cell lines expressed appreciable levels of ROR1 on the cell membrane, with the highest expression observed in mantle cell lymphoma (MCL) models compared with other B-cell lymphoma subtypes, and detectable expression in a few diffuse large B-cell lymphoma (DLBCL) and one marginal zone lymphoma (MZL) line. ROR1 RNA and surface protein levels were highly correlated. As a proof of principle, four ROR1-positive DLBCL cell lines and one cell line with low surface protein were exposed to the ROR1-targeting ADC zilovertamab vedotin for 120 hours. All ROR1-positive models were sensitive, with IC50 values ranging from 28 to 58 nM, whereas the remaining one was largely resistant. In summary, we defined a set of well-characterized lymphoma models suitable for preclinical studies of ROR1-directed agents, confirming the functional relevance of ROR1 surface expression for the activity of zilovertamab vedotin.

The gene coding for the receptor tyrosine kinase-like orphan receptor 1 (*ROR1*) was initially identified while looking for genes encoding proteins with tyrosine kinase-like domains ^1,2^. It is now understood that ROR1 possesses intrinsic tyrosine kinase activity and also functions as a receptor for Wnt5a, which activates β-catenin-independent non-canonical pathways, sustaining cell survival and oncogenesis ^2,3^. In adult tissues, including normal B lymphocytes, ROR1 is preferentially expressed by cancer cells rather than normal cells and has a role in sustaining the growth and survival of neoplastic cells ^2,4^. Several therapeutic strategies targeting ROR1 have been developed, including naked monoclonal antibodies, antibody-drug conjugates (ADCs), small molecule inhibitors, bispecific T cell engager (BiTE), and chimeric antigen receptor (CAR) T cells have been developed ^2^, with some currently in advanced clinical evaluation ^2^.

Here, we characterized a vast panel of lymphoma cell lines for their ROR1 RNA level and ROR1 cell surface protein expression, and showed, using zilovertamab vedotin as proof-of-principle, that these models can be exploited to study this promising class of anti-cancer agents.

## Material and methods

### Cell lines

Human lymphoma cell lines were cultured in the appropriate medium supplemented with fetal bovine serum (10% or 20%) and penicillin-streptomycin-neomycin (≈5,000 units penicillin, 5 mg streptomycin, and 10 mg neomycin/mL; Sigma-Aldrich, Darmstadt, Germany). Cell line identities were confirmed by short tandem repeat DNA fingerprinting using the Promega GenePrint 10 System kit (B9510). Cells were periodically tested for Mycoplasma negativity using the MycoAlert Mycoplasma Detection Kit (Lonza, Visp, Switzerland).

### Compounds

Zilovertamab vedotin was provided by Oncternal Therapeutics.

### Cytotoxic activity in single and combination

Cells were manually seeded in 96-well plates at a concentration of 50,000 cells/mL (10,000 cells in each well). Treatments were performed manually. After 120 hours of exposure to zilovertamab vedotin (11 concentrations, 1:2 dilutions starting from 100 nM) or DMSO, as control, cell viability was determined using 3-(4,5-dimethyl-thiazol-2-yl)-2,5-diphenyltetrazolium bromide (MTT), and the reaction was stopped after 4 hours with sodium dodecyl sulfate lysis buffer.

### ROR1 expression by flow cytometry

ROR1 expression was determined by flow cytometry on fresh cells. Cells were washed with ice-cold FACS buffer (PBS + 0.5% BSA) and divided 1x106 cells per tube. A pretreatment with human FcR blocking (Miltenyi Biotec Inc., Auburn, CA, USA) was performed according to manufacturer’s instructions. PE mouse anti-human ROR1 (clone 4A5) or PE mouse IgG2b, κ isotype control (clone 27-35) were incubated with cells at 4°C for 30 minutes; cells were washed twice in FACS buffer, and the final cell pellets were re-suspended in FACS buffer. Flow-cytometry analysis was carried out with a FACSCanto II instrument (BD Biosciences, Allschwil, Switzerland). Median Fluorescence intensity (MFI) of each sample was defined using FacsDiva v8.0.1 software (BD Biosciences, Allschwil, Switzerland). Unstained cells and cells stained with isotype control antibody were used as controls. Data were analyzed using FlowJo software version 11 (TreeStar Inc., Ashland, OR, USA).

### RNA expression

RNA expression values were extracted from the previously reported dataset obtained using total-RNA-Seq (GSE221770 ^5^).

### Data Analysis

Differences in expression among lymphoma subtypes were calculated using the non-parametric Mann-Whitney t-test. Statistical analyses, correlations, and boxplots were performed using Prism software v10.2.3 (GraphPad Software, La Jolla, CA, USA). A p-value < 0.05 was considered statistically significant.

## Results and discussion

To assess the expression levels of ROR1 across B-cell lymphoma cell lines, we first took advantage of an RNA-Seq dataset we had obtained on 47 B-cell lymphoma cell lines ^5^. ROR1 was expressed at the RNA level in most cell lines, although only 19 presented moderate to high expression (Figure 1). We then evaluated whether the ROR1 protein was expressed on the cell surface by flow cytometry in 29 cell lines (Supplementary Table S1). ROR1 was present on the surface of only a few cell lines (Figure 2A). Cell lines with high expression were mostly derived from mantle cell lymphoma (MCL) (median MCL =4.26; n=9, 95% C.I. 3.65-5.46) compared to all the other B-cell lymphoma cells tested (median others =0.87, n=20, 95% C.I. 0.43-1.47) (Figure 2B), a few diffuse large B-cell lymphoma (DLBCL) cell lines (median=0.87; n=16; 95% C.I. 0.4-1.65), and one marginal zone lymphoma (MZL) cell line (median=2.46; n=2; 95% C.I. 0.3-4.62) (Figure 2C). These findings are in line with what has been reported by others ^6-8^. RNA and protein levels were highly significantly correlated (r=0.82, P<0.0001) (Supplementary Figure 1).

**Figure 1.**
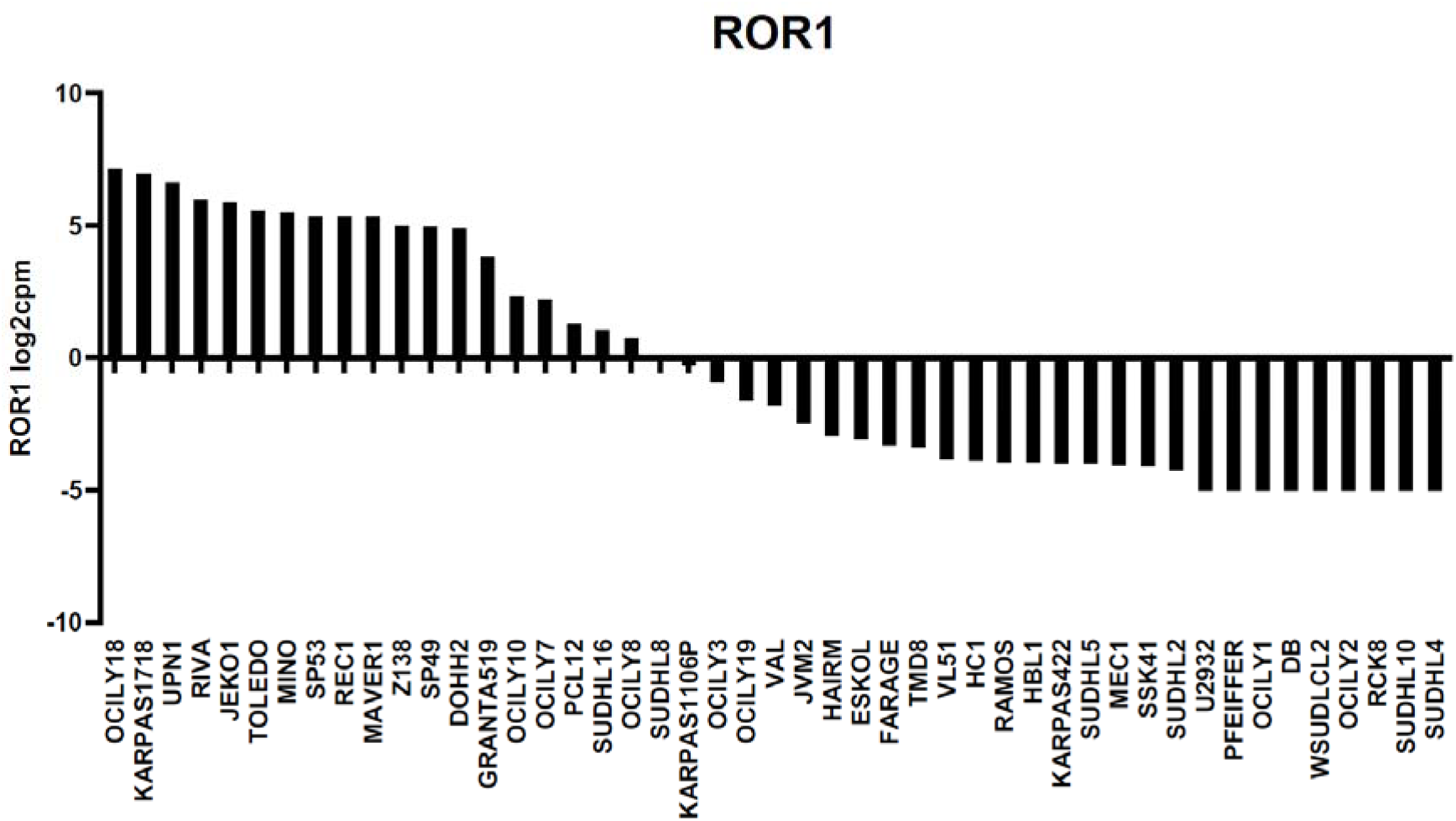
Variable ROR1 RNA expression across B cell lymphoma cell lines. X axes, lymphoma cell lines; Y axes, transcript expression measured by total RNA-Seq in log2cpm.

**Figure 2.**
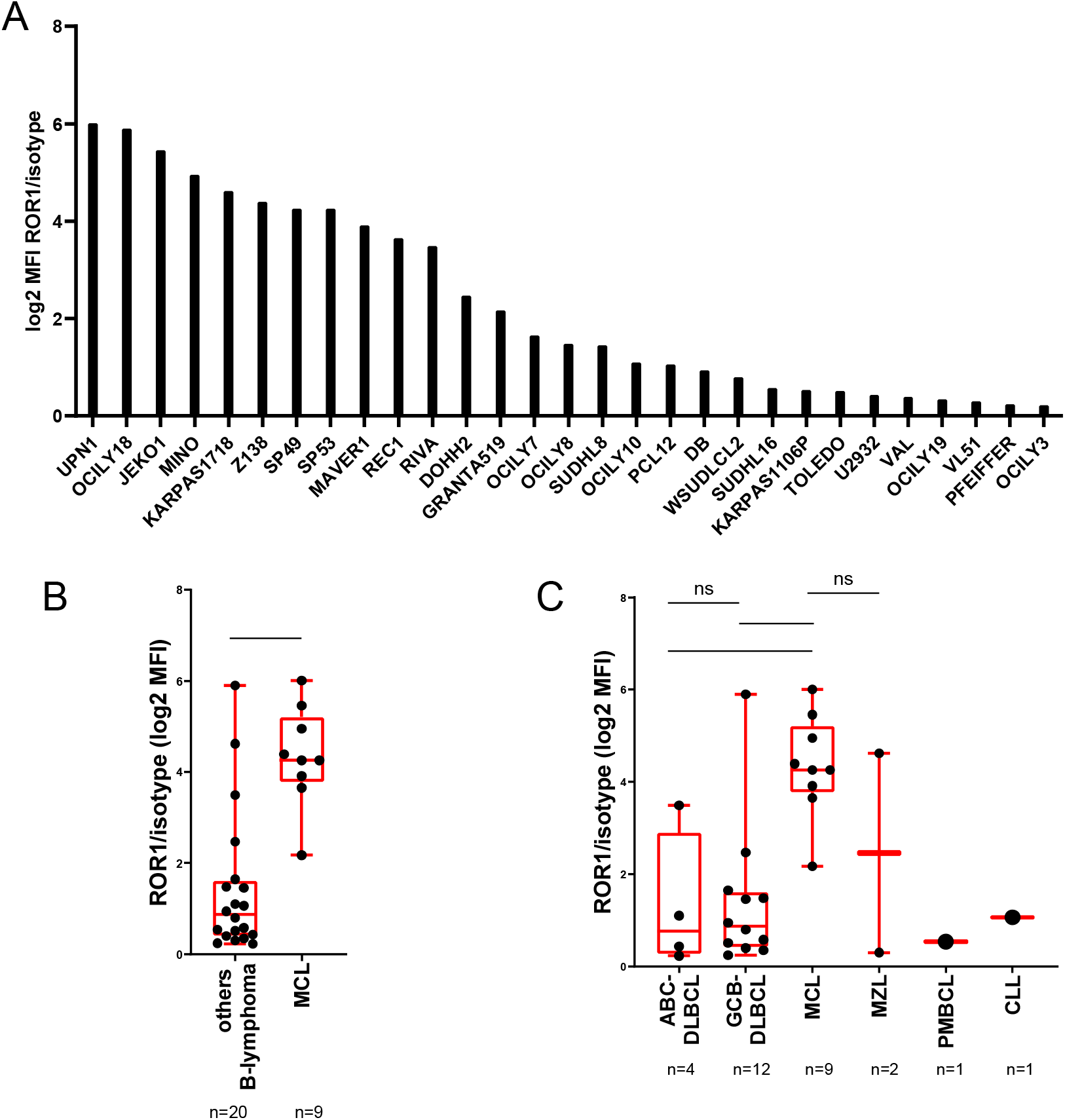
Variable cell surface protein expression across B-cell lymphoma cell lines. A) X axes, lymphoma cell lines; Y axes, protein expression measured by flow cytometry and calculated by log2 Median Fluorescence Intensity (MFI) of ROR1-isotype ratio. B) ROR1 expression in MCL and all the other lymphoma B-cell line subtypes. C) ROR1 expression among different subtypes of lymphoma. In each box plot, the line in the middle of the box represents the median, and the box extends from the 25th to the 75th percentile (interquartile range, IQ); the whiskers extend to the upper and lower adjacent values (i.e., ±1.5 IQ). Y-axis: log2 MFI values of ROR1-isotype ratio. DLBCL, diffuse large B-cell lymphoma; ABC, activated B cell; GCB, germinal center B cell; MCL, mantle cell lymphoma; MZL, marginal zone lymphoma; PMBCL, primary mediastinal large B-cell lymphoma; CLL, chronic lymphocytic leukemia. A non-parametric Mann-Whitney t-test was applied. * = p<0.05; **** = p<0.0001.

Based on these findings, we exposed four ROR1-positive DLBCL models (DOHH2, RIVA, OCI-LY18, OCI-LY7) and the TOLEDO cell line, which had very low ROR1 protein levels despite having high RNA expression, to the ROR1-targeting zilovertamab vedotin (UC961-vc-MMAE) for 5 days. All the positive cell lines were sensitive, with IC50 values ranging from 28 nM to 58 nM. Conversely, zilovertamab vedotin did not show a significant activity in the TOLEDO cell line (Supplementary Figure 2; Supplementary Table S2).

In conclusion, we showed that ROR1 is expressed in several lymphoma cell lines at RNA and protein levels on the cell surface, with a high correlation between RNA and cell surface protein expression.

These models can be used to study ROR1-targeting agents, as shown using the ROR1-targeting ADC zilovertamab vedotin.

## Acknowledgements

This project was partially supported by research funds from Oncternal Therapeutics.

## Author Contributions

CT: performed experiments, performed data mining, interpreted data, and co-wrote the manuscript. EC, FS, GR: performed experiments, interpreted data.

LC: performed data mining.

GF: co-designed the study, provided reagents, and supervised the study.

FB: co-designed the study, interpreted data, supervised the study, and co-wrote the manuscript. All authors reviewed and accepted the final version of the manuscript.

## Potential conflicts of interest

CT: travel grant from iOnctura.

LC: institutional research funds from Orion; travel grant from HTG.

GK: Oncternal Therapeutics: employment.

FB: institutional research funds from ADC Therapeutics, Bayer AG, BeiGene, Floratek Pharma, Helsinn, HTG Molecular Diagnostics, Ideogen AG, Idorsia Pharmaceuticals Ltd., Immagene, ImmunoGen, Menarini Ricerche, Nordic Nanovector ASA, Oncternal Therapeutics, Spexis AG; consultancy fee from BIMINI Biotech, Floratek Pharma, Helsinn, Immagene, Menarini, Vrise Therapeutics; advisory board fees to institution from Novartis; expert statements provided to HTG Molecular Diagnostics; travel grants from Amgen, AstraZeneca, iOnctura.

The other authors have no conflicts of interest.

## References

1. Masiakowski P, Carroll RD. A novel family of cell surface receptors with tyrosine kinase-like domain. J Biol Chem. 1992;267(36):26181–26190.

2. Tigu AB, Munteanu R, Moldovan C, et al. Therapeutic advances in the targeting of ROR1 in hematological cancers. Cell Death Discov. 2024;10(1): 471.

3. Zhao Y, Zhang D, Guo Y, et al. Tyrosine Kinase ROR1 as a Target for Anti-Cancer Therapies. Front Oncol. 2021;11.

4. Kipps TJ. ROR1: an orphan becomes apparent. Blood. 2022;140(14): 1583–1591.

5. Johnson Z, Tarantelli C, Civanelli E, et al. IOA-244 is a Non-ATP-competitive, Highly Selective, Tolerable PI3K Delta Inhibitor That Targets Solid Tumors and Breaks Immune Tolerance. Cancer Res Commun. 2023;3(4):576–591.

6. Zhang Q, Wang HY, Liu X, et al. Cutting Edge: ROR1/CD19 Receptor Complex Promotes Growth of Mantle Cell Lymphoma Cells Independently of the B Cell Receptor-BTK Signaling Pathway. J Immunol. 2019;203(8): 2043–2048.

7. Jiang VC, Liu Y, Jordan A, et al. The antibody drug conjugate VLS-101 targeting ROR1 is effective in CAR T-resistant mantle cell lymphoma. J Hematol Oncol. 2021;14(1): 132.

8. Ghaderi A, Daneshmanesh AH, Moshfegh A, et al. ROR1 Is Expressed in Diffuse Large B-Cell Lymphoma (DLBCL) and a Small Molecule Inhibitor of ROR1 (KAN0441571C) Induced Apoptosis of Lymphoma Cells. Biomedicines. 2020;8(6).

