## Supplementary tables and figures for "Characterization of lymphoma models for the surface ROR1 expression"

### Supplementary figures and tables

**Supplementary Figure 1. ROR1 RNA levels and protein cell surface expression are correlated in B-cell lymphoma cell lines.** Linear regression between ROR1 transcript and protein expression is shown in black. Orange dots: GCB-DLBCL cell lines; blue dots: ABC-DLBCL cell lines; green dots: MCL cell lines; purple dots: MZL cell lines; grey dots: CLL cell lines; and black dots: PMBCL cell lines. X axis, ROR1 transcript expression as log2 cpm. Y axis, log2 MFI values of ROR1-isotype ratio. DLBCL, diffuse large B-cell lymphoma; ABC, activated B cell; GCB, germinal center B cell; MCL, mantle cell lymphoma; MZL, marginal zone lymphoma; PMBCL, primary mediastinal large B-cell lymphoma; CLL, chronic lymphocytic leukemia. Two-tailed Pearson correlation was calculated.

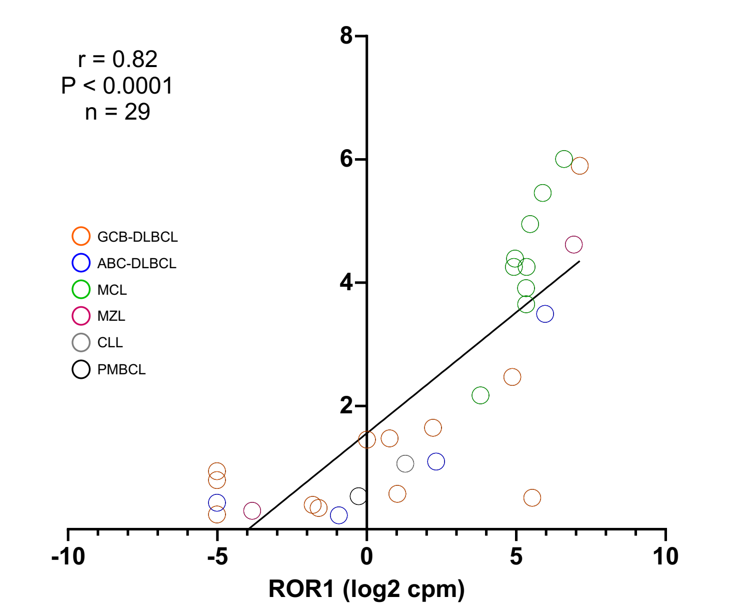


**Supplementary Figure 2. Assessment of the anti-proliferative activity of zilovertamab vedotin in lymphoma cell lines after 120 hours of exposure.** Five DLBCL cell lines have been exposed to increasing concentrations of zilovertamab vedotin for five days. X-axis: increasing concentrations of zilovertamab vedotin in nM. Y-axis: cell viability in %. DLBCL, diffuse large B-cell lymphoma.

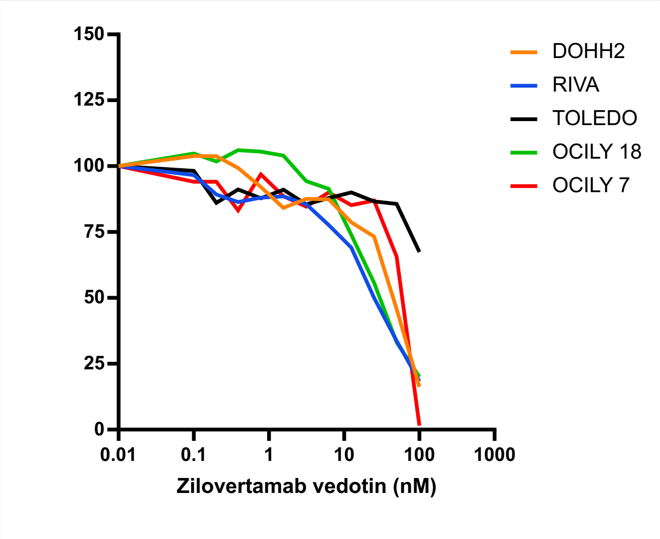

**Supplementary Table 1. Surface ROR1 protein expression on the cell surface as detected by flow cytometry.**

| Cell line | Histology | MFI ROR1/isotype |
|---|---|---|
| OCI-LY10 | ABC-DLBCL | 2.14 |
| OCI-LY3 | ABC-DLBCL | 1.17 |
| RIVA | ABC-DLBCL | 11.26 |
| U2932 | ABC-DLBCL | 1.35 |
| PCL12 | CLL | 2.09 |
| DB | GCB-DLBCL | 1.92 |
| DOHH2 | GCB-DLBCL | 5.54 |
| OCI-LY18 | GCB-DLBCL | 59.60 |
| OCI-LY19 | GCB-DLBCL | 1.27 |
| OCI-LY7 | GCB-DLBCL | 3.13 |
| OCI-LY8 | GCB-DLBCL | 2.78 |
| PFEIFFER | GCB-DLBCL | 1.18 |
| SU-DHL-16 | GCB-DLBCL | 1.49 |
| SU-DHL-8 | GCB-DLBCL | 2.74 |
| TOLEDO | GCB-DLBCL | 1.42 |
| VAL | GCB-DLBCL | 1.32 |
| WSUDLCL2 | GCB-DLBCL | 1.74 |
| GRANTA519 | MCL | 4.51 |
| JEKO1 | MCL | 43.89 |
| MAVER1 | MCL | 15.05 |
| MINO | MCL | 30.92 |
| REC1 | MCL | 12.56 |
| SP49 | MCL | 19.11 |
| SP53 | MCL | 19.10 |
| UPN1 | MCL | 64.33 |
| Z138 | MCL | 20.98 |
| KARPAS1718 | MZL | 24.56 |
| VL51 | MZL | 1.23 |
| KARPAS1106P | PMBCL | 1.45 |

DLBCL, diffuse large B-cell lymphoma; ABC, activated B cell; GCB, germinal center B cell; MCL, mantle cell lymphoma; MZL, marginal zone lymphoma; PMBCL, primary mediastinal large B-cell lymphoma; CLL, chronic lymphocytic leukemia.

**Supplementary Table 2. IC50 values obtained by exposing DLBCL cell lines to zilovertamab vedotin for 120 hours.**

| cell line | IC50 zilovertamab vedotin (nM) |
|---|---|
| DOHH2 | 44.08 |
| RIVA | 29.61 |
| TOLEDO | 167.4 |
| OCI-LY18 | 27.69 |
| OCI-LY7 | 57.61 |
